# Anatomically remote education of B cells is required for colonic health

**DOI:** 10.1101/806497

**Authors:** Neeraj K. Surana, Cheryn J. Couter, David Alvarez, Uli H. von Andrian, Dennis L. Kasper

**Affiliations:** Department of Immunology, Harvard Medical School, Boston, MA, USA; Division of Infectious Diseases, Department of Medicine, Boston Children’s Hospital, Boston, MA, USA; The Ragon Institute of Massachusetts General Hospital, Massachusetts Institute of Technology and Harvard University, Cambridge, MA, USA; Departments of Pediatrics, Molecular Genetics and Microbiology, and Immunology, Duke University, Durham, NC, USA

## Abstract

Mucosa-associated lymphoid tissues contain roughly 80% of all immune cells and produce virtually all of the body’s IgA^1–3^. Although the majority of IgA-secreting cells educated within a mucosal site home back to the same anatomic region, some cells are also found in distant mucosal tissues^2–6^. These observations underlie the notion of a common mucosal immune system, which holds that anatomically unrelated mucosal sites are functionally connected by a shared immune system^2,3^. However, the ontological basis of this separation between site of immune education and functionality has remained elusive. Here we show that mice lacking Peyer’s patches (PPs)—small-intestinal lymphoid tissue covered by antigen-sampling M cells—have no immunologic defect in the small-intestinal lamina propria. Surprisingly, the primary immunological abnormality in PP-deficient mice was a reduction in colonic B cells, including plasmablasts but not plasma cells. Adoptive transfer experiments conclusively demonstrated that PP-derived cells preferentially give rise to colonic—but not small-intestinal—B cells and plasmablasts. Finally, these PP-derived colonic B cells were critical for restraining colonic inflammation. Thus, PPs bridge the small-intestinal and colonic immune systems and provide a clear example of immune education being required in an anatomic compartment distinct from the effector site. Our findings, which highlight that the majority of fecal IgA is produced by colonic plasmablasts that originate from PPs, will help inform design of mucosal vaccines.

## Main Text

It has long been known that the mucosal immune system is functionally interrelated, with antigens delivered to one mucosal site evoking an immune response in remote mucosal tissues^2,3,7^. Although mucosal lymphocytes have an anatomic affinity for the mucosa of origin, these cells and their antibodies can be found throughout mucosa-associated lymphoid tissues and, to a lesser extent, systemic compartments^2–6^. While it is known that there is some compartmentalization within this common mucosal immune system^2,3^, it remains unclear whether this anatomic separation between site of education and functionality is essential to maintaining immune homeostasis.

Given that Peyer’s patches (PPs)—small-intestinal subepithelial lymphoid aggregates that are covered by a follicle-associated epithelium, including M cells that are specialized for the uptake and delivery of luminal antigens to underlying immune cells—are a primary site for immune education within the small intestine^1^, we explored how PPs influence the immune system. Pregnant mice were injected at embryonic day 14 (E14) or E15 with an antibody to interleukin (IL) 7Rα, thereby preventing formation of PPs in their pups^8^. Unlike genetically deficient animals that lack PPs, this approach involves only a transient depletion of lymphocytes during a critical window in embryogenesis and has been demonstrated not to affect formation of small-intestinal isolated lymphoid follicles, the cecal patch, or mesenteric lymph nodes (MLNs)^8,9^. We found that colonic patches are similarly unaffected (data not shown).

Given that M cell formation is dependent on signals from PPs^1,10^, we reasoned that PP-deficient mice also lack M cells. Consistent with this hypothesis, we found that PP-deficient mice have fewer viable commensal organisms in the MLNs that drain the small intestine than do control mice (Fig. 1A); this observation highlighted the critical role M cells play in transcytosing small-intestinal bacteria^1,11^. Furthermore, we found that PP-deficient mice orally infected with *Salmonella enterica* serotype Typhimurium—a pathogen known to adhere to and invade via M cells^12^—have a lower small-intestinal burden of mucosa-associated *Salmonella* than do control mice (Fig. 1B), a result further suggesting reduced M cell number and/or function. Indeed, flow cytometric analysis of small-intestinal epithelial cells revealed reduced numbers of GP2^+^ cells, a marker for M cells^13^, in PP-deficient mice (Fig. 1C). Moreover, examination of the distal ileum of PP-deficient mice by transmission electron microscopy demonstrated a complete lack of M cells (Fig. 1D). Finally, consistent with the notion that PPs are a critical inductive site for intestinal immunity, PP-deficient mice have drastically reduced levels of fecal IgA (Fig. 1E). Thus, mice treated with an antibody to IL-7Rα at E14 or E15 have a specific loss of Peyer’s patches and M cells, with a resulting defect in IgA production.

**Figure 1.**
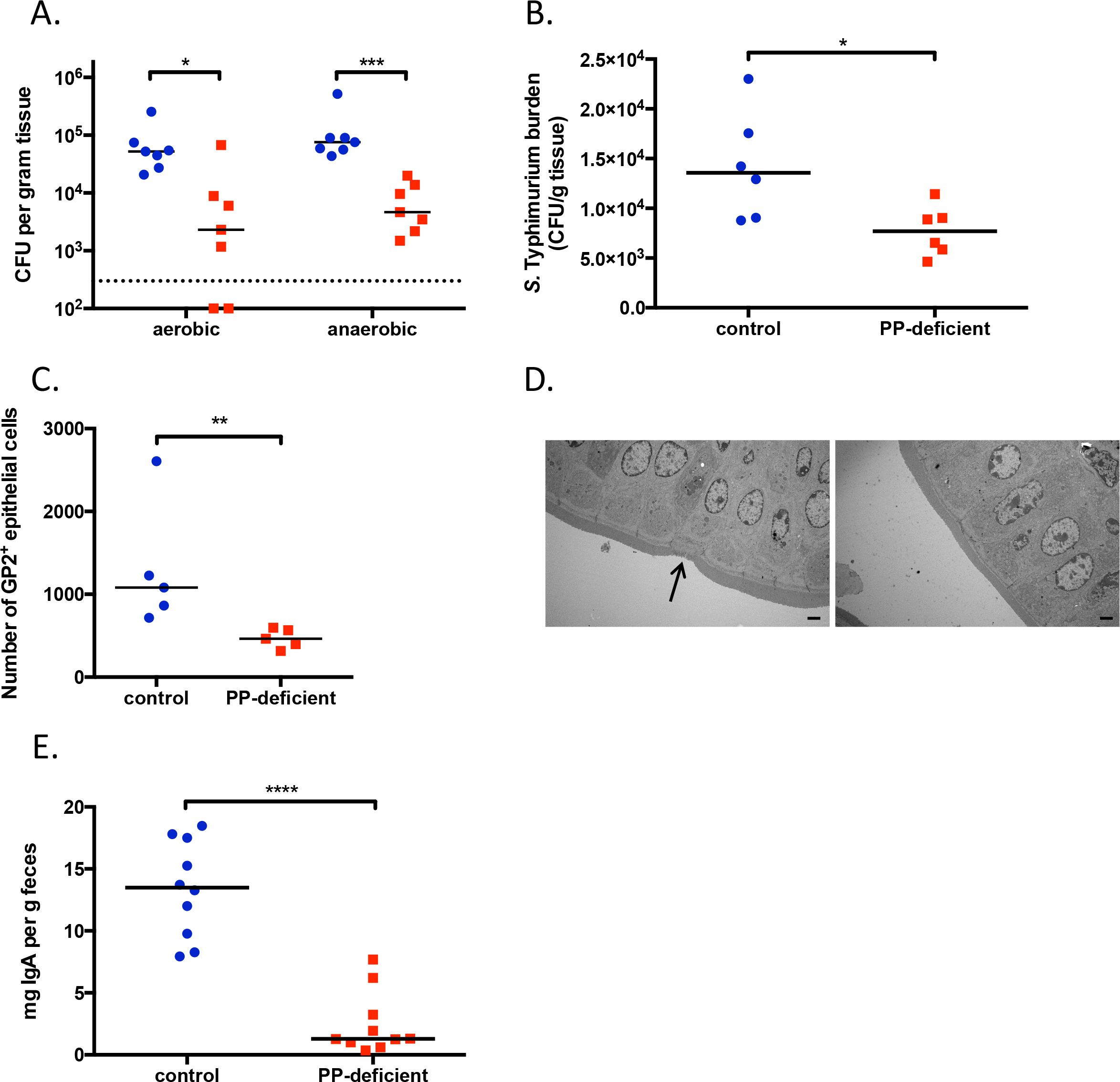
PP-deficient mice lack M cells and have impaired production of IgA. A. CFU of aerobic and anaerobic bacteria cultured from MLNs. The dotted line represents the limit of detection. B. Ileal burden of mucosa-associated *S*. Typhimurium after oral gavage of 10^8^ CFU. C. Number of GP2^+^ cells in the small-intestinal epithelial layer. D. Transmission electron microscopy of the distal ileum. Arrow in left panel points to an M cell. Images are magnified 1500x, and the bar represents 2 μm. E. Levels of fecal IgA. For all figures, control animals are shown in blue circles, PP-deficient animals are depicted by red squares. NS, not significant; *, *p*<0.05; **, *p*<0.01; ***, *p*<0.001; ****, *p*<0.0001. Data are representative of ≥2 independent experiments.

To more fully understand the impact of PPs on development of the immune system, we performed flow cytometric analyses of various immune compartments. Consistent with previous reports, the loss of PPs has no impact on the number of CD45^+^ lymphocytes in the spleen, MLNs, or small-intestinal lamina propria (LP) (Fig. 2A)^8,14,15^. Thus, PPs are dispensable for the development of these immune organs. Surprisingly, PP-deficient mice have a lymphopenic colon (Fig. 2A). Further characterization of these cells suggested no change in the numbers of colonic TCRβ^+^ cells, TCRγδ^+^ cells, or various myeloid cell populations (Fig. 2B, Extended Data Fig 1); rather, there is a significant decrease in the frequency and number of colonic CD19^+^ cells (Fig. 2C, 2D). The fact that this decrease in colonic B cells accounted for more than two-thirds of the decrease in colonic CD45^+^ cells indicated that this is the primary cell population lost in the colons of PP-deficient mice. This unexpected finding raised the question of whether PPs are important for the accumulation of B cells in other mucosal sites. To address this question, we examined the lungs of PP-deficient mice and found that they have a normal frequency of B cells (Fig. 2E). Given that PPs are known to be a central site for the development of IgA-secreting B cells, we assessed whether IgA^+^ B cells in the colon of PP-deficient mice are similarly affected. Intriguingly, PP-deficient mice have a decreased frequency and number of colonic B220^+^IgA^+^ cells (plasmablasts) but no defect in colonic B220^−^IgA^+^ cells (plasma cells) (Fig. 2F, Extended Data Fig. 2A). In contrast, there is no difference from control mice in small-intestinal plasmablasts or plasma cells (Extended Data Fig. 2B). In summary, we have established that the absence of small-intestinal PPs affects the ontogeny of B cells, including plasmablasts, specifically in the colon—an anatomically remote site.

**Figure 2.**
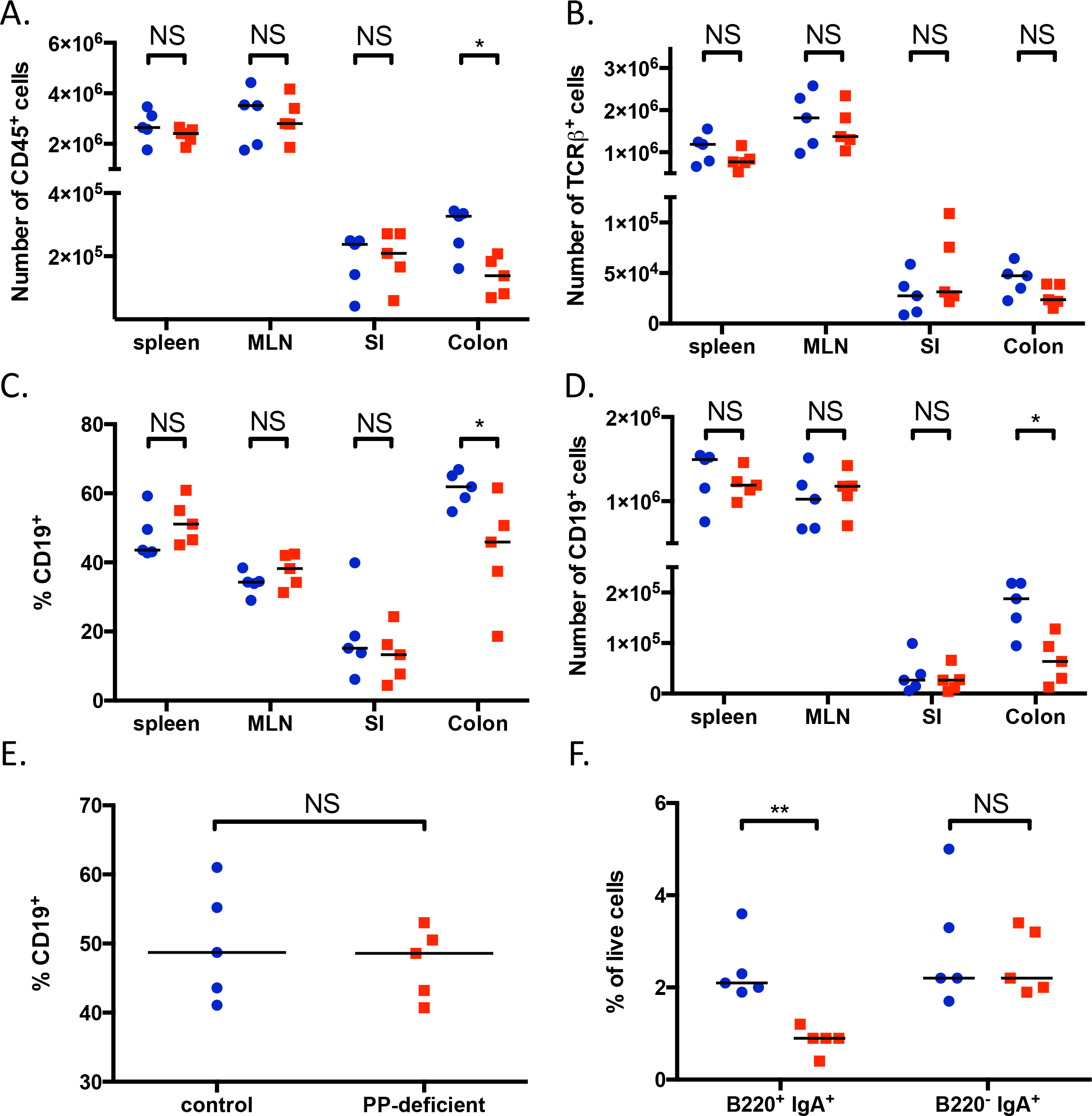
PP-deficient mice have decreased numbers of colonic B cells. A, B, D. Numbers of CD45^+^ (A), TCRb^+^ (B), and CD19^+^ (D) cells in the spleen, MLNs, small-intestinal lamina propria (SI), and colonic lamina propria. C. Frequencies of CD19^+^ cells (gated on live CD45^+^ cells) in the same organs. E. Frequency of CD19^+^ cells in the lung. F. Frequencies of plasmablasts (B220^+^ IgA^+^) and plasma cells (B220^−^ IgA^+^) in the colonic lamina propria. NS, not significant; *, p<0.05; **, p<0.01. Data are pooled froma minimumof 3 independent experiments.

It is not clear how the loss of PPs leads to a decrease in colonic B cells. One possibility is that prenatal treatment with IL-7Rα antibody affects—in addition to PP development—some other potential source of colonic B cells. It has been demonstrated that peritoneal B1 cells can accumulate in the small-intestinal LP, ultimately developing into plasma cells^16^. It is possible that PP-deficient mice also have a defect in a population of peritoneal B cells that ultimately traffics to the colon. Analyzing the number of CD19^+^ cells in peritoneal lavage fluid, we found no difference between control and PP-deficient mice (Fig. 3A). Moreover, PP-deficient mice have a normal number of colonic B1 cells (CD19^+^ CD43^+^), B1a cells (CD19^+^ CD43^+^ CD5^+^), B1b cells (CD19^+^ CD43^+^ CD5^−^), and regulatory B cells (CD19^+^ CD43^+^ CD5^+^ CD1d^+^) (Extended Data Fig. 3 and data not shown); these results indicate that PP-deficient mice lack colonic B2 cells. In addition, it has recently been reported that some colonic IgA-secreting cells originate from the cecal patch^17^; however, the normal number of cecal B cells in PP-deficient mice (Fig. 3B) suggests that the decrease in colonic B cells is not due to altered numbers of B cells in the cecum.

**Figure 3.**
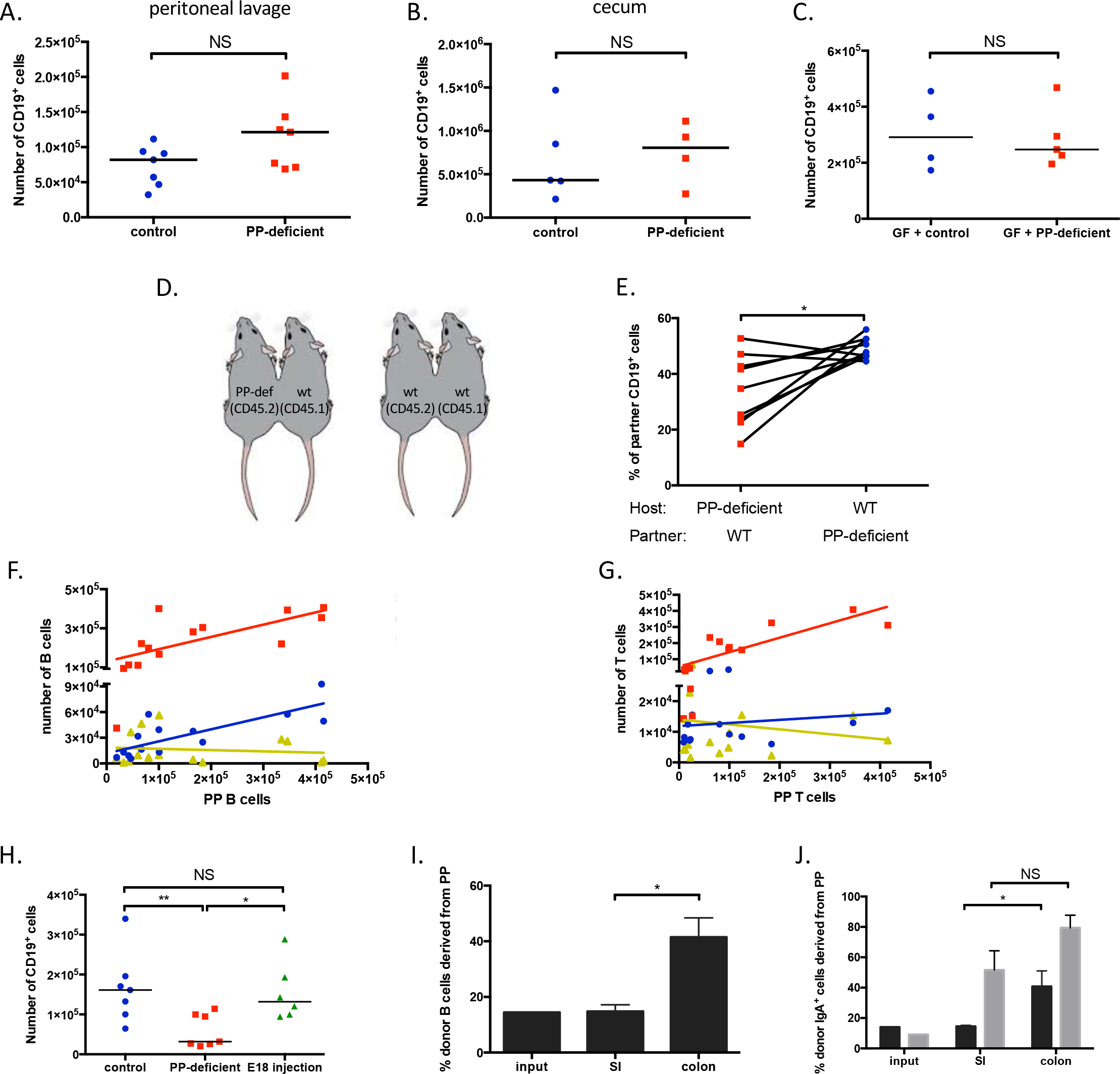
A subset of colonic B cells is derived from Peyer’s patches. A, B. Numbers of CD19^+^ cells in peritoneal lavage fluid (A) and the cecum (B). C. Number of CD19^+^ cells in the colonic lamina propria of GF animals colonized with the microbiota of either control or PP-deficient mice. D. Schematic of parabiotic mice generated. E. Percentage of colonic CD19^+^ cells in the indicated host mouse that is derived from the parabiont partner. Lines connect parabiont partners. F, G. Correlation of the numbers of CD19^+^ (F) and TCRβ^+^ (G) cells between PPs and the spleen (red squares), colon (blue circles), or small intestine (SI; yellow triangles). H. Number of colonic CD19^+^ cells in control mice, PP-deficient mice (exposed to an antibody to IL-7Rα at E15), and mice prenatally exposed to IL-7Rα antibody at E18. I, J. Fraction of PP-derived CD19^+^ (I) and IgA^+^ (CD19^+^ in dark bars and CD19^−^ in gray bars) (J) cells in the input, the SI, or the colon (n=4 recipients for I and J). NS, not significant; *, p<0.05; **, p<0.01. Data are pooled (A, B, E.H) or representative of ≥2 independent experiments (C, I, J).

Given the growing body of evidence on the microbiota’s impact on the development of the immune system^7,18^, we wondered whether *in utero* antibody treatment had affected the microbiota of the dams—and ultimately that of their pups—in a manner that specifically affects colonic B cells. To test this hypothesis, we transferred the fecal microbiota of control or PP-deficient mice into germ-free (GF) mice. Three weeks later, we assessed the frequency and number of colonic B cells in these mice. The lack of difference (Fig. 3C, Extended Data Fig. 4) proves that the colonic B-cell defect in PP-deficient mice is not due to an altered microbiota.

Finally, we considered the possibility that prenatal antibody treatment had affected B-cell trafficking—either the ability of B cells to home to the colon or the capacity of the colon to recruit B cells. Analysis of colonic levels of MADCAM1 (a receptor that enables α4β7 integrin-expressing lymphocytes to home to the intestinal LP), CXCL13 (a chemokine that is selectively chemotactic for B cells), and CCL25 and CCL28 (chemokines relevant for homing of B cells to the small and large intestines, respectively) revealed no difference between control and PP-deficient mice (Extended Data Fig. 5), demonstrating that these trafficking molecules are not the cause of the colonic B-cell defect in PP-deficient mice. To more robustly examine B-cell trafficking, we performed parabiotic experiments using mice that were congenic for CD45 allotypes (i.e., CD45.1 and CD45.2) and were either PP-deficient or PP-sufficient (wild-type, WT) (Fig. 3D). As a control, we generated parabiotic pairs of congenic WT mice. As expected, expression of different CD45 allotypes does not affect recruitment of B or T cells in the spleen, small intestine, or colon (Extended Data Fig. 6). While PP-deficient mice have no defect in the recruitment of WT B or T cells in the spleen (Extended Data Fig. 7), there is a defect in the recruitment of B cells to the colon (Fig. 3E). Given that PP-deficient mice also have a defect in recruitment of B cells to the small intestine and of T cells to the colon and small intestine (without an alteration of cell numbers) (Extended Data Fig. 7), it is not clear that the recruitment defect of colonic B cells fully explains the decreased number of colonic B cells in PP-deficient mice.

An alternative explanation for the paucity of colonic B cells in PP-deficient mice is that a portion of colonic B cells are derived from PPs with no redundant source. If so, we reasoned that the number of colonic B cells should be correlated to that of PPs. Using WT (i.e., PP-sufficient) mice ranging in age from 1 to 18 weeks, we compared the number of B cells in the colon or small intestine with that in PPs. Strikingly, we found a strong correlation in B cell numbers between the colon and PPs (Spearman correlation 0.77; *p*=0.002) but not between the small intestine and PPs (Spearman correlation 0.00; *p*=0.99) (Fig. 3F). Of note, we remove PPs from the small intestine prior to analysis, but we are technically unable to excise colonic patches before analyzing the colon. If the correlation observed between colonic and PP B cells was due simply to the residual presence of a lymphoid structure in the colon, we predict that numbers of both B and T cells would be correlated given their shared trafficking route through lymphatics. As a control, we analyzed splenic lymphocyte numbers and found a robust correlation of both B and T cells in the spleen and PPs (Fig. 3F, 3G). However, there is no correlation between T cells in PPs and either the colon or small intestine (Fig. 3G). These results indicate that the colon-PP correlation that is limited to B cells is due to something other than the presence of colonic patches and suggest a direct link between these B-cell populations.

Next, we exploited the fact that administration of the IL-7Rα antibody at different gestational ages leads to a different number of PPs to test whether the number of PPs present in a mouse is similarly correlated to the number of colonic B cells (Extended Data Fig 8A)^8^. As in the previous experiment relying on mice of various ages, we found—using age-matched mice that differed only in the number of PPs—that the number of colonic B cells is tightly linked to the number of PPs (Extended Data Fig. 8B; Spearman correlation 0.80; *p*=0.0009). Analysis of mice that were exposed to the IL-7Rα antibody at E18 and had a normal number of PPs revealed a number of colonic B cells similar to that in control mice and greater than that in PP-deficient animals (Fig. 3H). Taken together, these data effectively limit any potential off-target effect of the antibody to another structure that, similar to PPs, increases in number between E15 and E18 in an IL-7Rα-dependent, stepwise fashion. Moreover, these correlations significantly strengthen the notion that PP-derived B cells form a distinct population of colonic B cells

To more conclusively determine whether a subset of colonic B cells is derived from PPs, we performed adoptive transfer experiments, injecting a mixture of PP cells (CD45.1^+^) and a 6-fold excess of splenocytes (CD45.1^+^ CD45.2^+^) into PP-deficient mice (CD45.2^+^). If PPs truly represent a non-redundant source for a subpopulation of colonic B cells, we conjectured that the donor-derived B cells in the colon would contain a greater percentage of PP cells than the injected mixture. Indeed, while the small intestine contains the same fraction of PP-derived donor B cells as the input, significantly more PP-derived donor B cells are present in the colon of recipient mice (Fig. 3I). In contrast, the frequency of PP-derived T cells in the colon does not increase (Extended Data Fig. 9). When we analyzed IgA^+^ B cells in these mice, we found that PP-derived cells preferentially give rise to plasma cells (CD19^−^ IgA^+^) in the small intestine (Fig. 3J); this result was similar to that obtained by Craig and Cebra in seminal work^6^. Extending this work, we found significantly more PP-derived plasma cells in the colon as well. Interestingly, the colon—but not the small intestine—also contains significantly increased numbers of PP-derived plasmablasts (CD19^+^ IgA^+^). These data confirm that PP-derived B cells, including plasmablasts, preferentially home to the colon to a greater degree than splenic B cells. Taken together, our data establish the existence of a cohort of colonic B cells that originates from PPs.

To determine whether these PP-derived colonic B cells are functionally important, we subjected mice to infection with *Citrobacter rodentium*, a model of enteropathogenic and enterohemorrhagic *Escherichia coli* infections in whose control B cells are central^19,20^. PP-deficient mice have fecal and colonic burdens of *C. rodentium* greater than those in control mice (Fig. 4A, 4B). Thus, the loss of PP-derived colonic B cells results in an inability to control infectious colitis.

**Figure 4.**
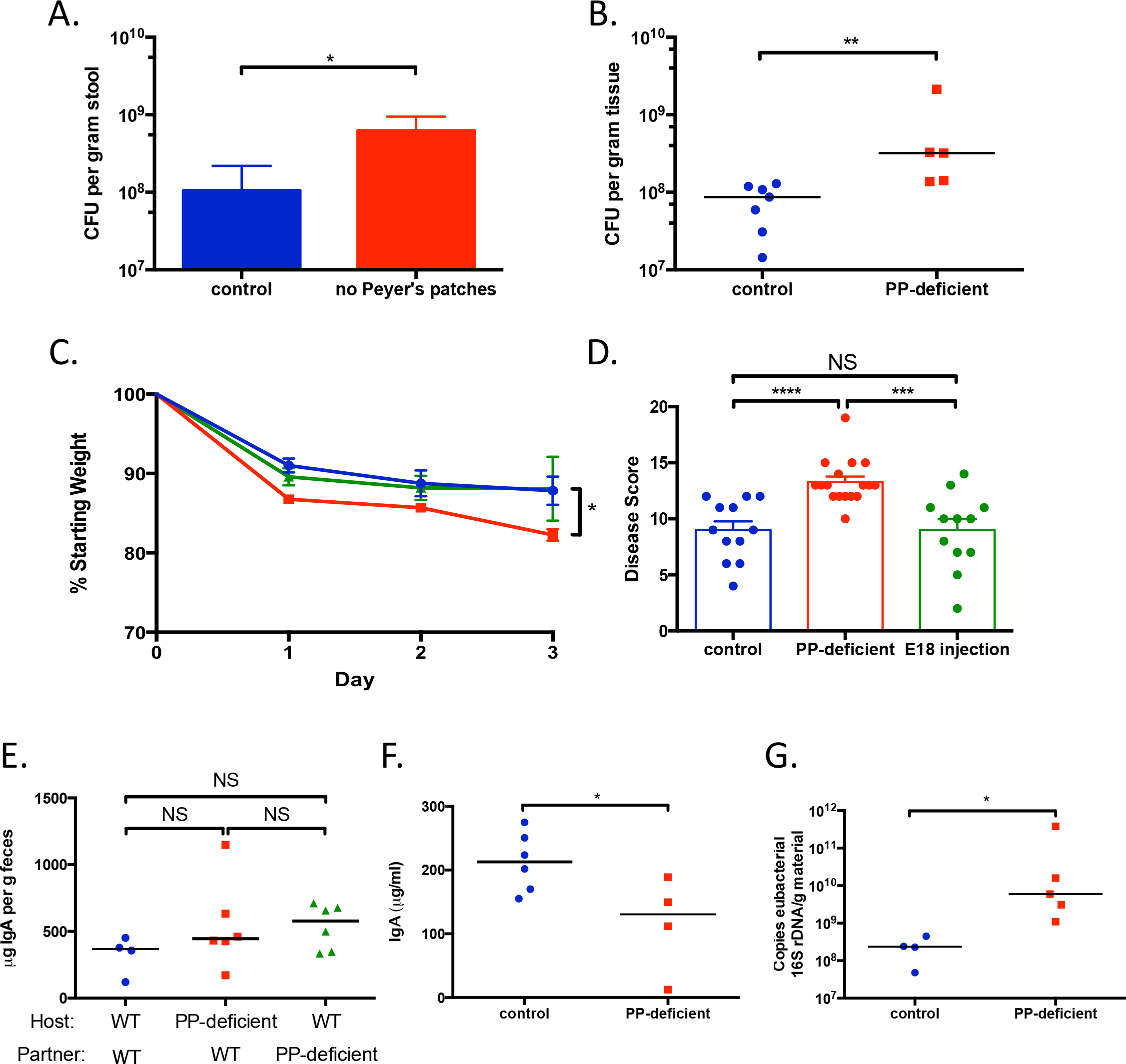
PP-derived colonic B cells regulate the colonic immune response in inflammatory and healthy conditions. A, B. Fecal (A) and colonic (B) burden of *C. rodentium* 8 days after oral gavage of control and PP-deficient mice with 5 × 10^8^ CFU of *C. rodentium*. C, D. Serial weights (C) and disease score (D) for control mice (blue circles) and mice prenatally exposed to IL-7Rα antibody at E14 (PP-deficient; red squares) or E18 (green triangles) and then subjected to TNBS-induced colitis. E. Levels of fecal IgA in the indicated host mouse 2 weeks after parabiosis with the listed partner. F. Colonic segments were harvested from control and PP-deficient mice and cultured *ex vivo*. The IgA concentration in the supernatant was measured. G. qPCR-based quantification of mucosa-associated bacteria in the colon. NS, not significant; *, *p*<0.05; **, *p*<0.01; ***, *p*<0.001; ****, *p*<0.0001. Data are pooled (C-D) or representative of ≥2 independent experiments (A–B, E–G).

Multiple studies have established that a subset of patients with Crohn’s disease—particularly those with more severe disease—have elevated titers of autoantibodies to GP2^21–23^, a receptor present on M cells^13^. Therefore, it has been speculated that M cells and PPs may play a role in the pathogenesis of Crohn’s disease^24,25^. To probe this hypothesis and assess whether PP-derived colonic B cells might specifically be involved, we exposed mice to trinitrobenzensulfonic acid (TNBS) in order to induce colitis in a murine model of Crohn’s disease^26^. PP-deficient animals lose more weight and have higher disease scores than control mice (Fig. 4C, 4D). Moreover, animals that are exposed to the IL-7Rα antibody at E18 and have normal numbers of PPs and colonic B cells display disease activity similar to that in control animals and less severe than that in PP-deficient animals (Fig. 4D). These findings linking absent M cell and PP function with more severe colonic inflammation may provide a mechanistic explanation for the association of antibodies to GP2, which likely alter M-cell function, with more severe Crohn’s disease. Taken together, these results establish that PP-derived colonic B cells are indispensable in controlling colonic health in both infectious and chemically induced forms of colitis.

The experiments reported herein demonstrate that PP-derived colonic B cells play a dramatic role in restraining colonic inflammation. We questioned whether these B cells regulate immune responses under healthy, homeostatic conditions. Although PP-deficient mice have dramatically lower levels of fecal IgA than do control mice (Fig. 1E), parabiosis to a PP-sufficient mouse, which allows migration of PP-derived B cells into the PP-deficient mouse, can restore fecal IgA levels (Fig. 4E). To examine regional differences in the production of IgA, we measured IgA levels in distal ileal contents as a measure of small-intestinal IgA. Surprisingly, PP-deficient mice have levels of small-intestinal IgA similar to those in control mice (Extended Data Fig. 10A)—an indication that the lack of PPs and the associated decrease in colonic plasmablasts result in a loss of IgA produced in the colon. Consistent with this finding, colonic segments from PP-deficient mice cultured *ex vivo* produce lower levels of IgA than those from control mice, whereas there is no difference between cultured ileal segments from the two groups (Fig. 4F, Extended Data Fig. 10B). These data support the finding that PPs are critical for IgA production and clarify that the bulk of fecal IgA originates in the colon, a finding consistent with human data^27^. Moreover, although counts of small-intestinal mucosal bacteria do not differ in the two groups of mice (Extended Data Fig. 10C), this defect in colon-specific IgA results in an increased colonic burden of mucosal bacteria in PP-deficient mice (Fig. 4G), which is associated with the development of colitis in mice and humans^28,29^. Thus, PP-derived colonic B cells are crucial regulators of the colonic immune response in both healthy and inflammatory conditions.

Our findings provide a conclusive example of homeostatic antigen sampling in one mucosal compartment being required for the ontogeny and functionality of cells in a distant site. From a teleological perspective, it is not readily apparent why an anatomic defect in the small intestine would lead to an altered immune status specifically in the colon. However, the fundamental physiologic roles of the small and large intestines—namely, the small intestine samples and absorbs luminal contents (e.g., nutrients), and, except for regulating water homeostasis, the large intestine largely allows unfettered passage of luminal contents—may partially explain why small-intestinal sampling of antigen is important for the colonic immune response. The anatomic characteristics of PPs—increased interactions with antigens facilitated by the presence of little or no overlying mucus, the presence of M cells that actively sample luminal antigens, and the fact that PPs are more numerous than corresponding structures in the colon—may underlie their indispensable role in the ontogeny of colonic B cells. Drawing parallels to the thymus—a specialized lymphoid organ that facilitates maturation of T cells—we suggest that PPs may be similarly specialized lymphoid organs that facilitate education of colonic B cells. While isolated lymphoid follicles and/or other mucosal sources may be able to compensate for small-intestinal B cells, the lack of other redundant sources for colonic B cells makes PPs a critical immunological sentinel for the colon.

## Acknowledgements

We thank Shakir Edwards for assistance with gnotobiotic mice; Roderick Bronson for review of histology; Julie McCoy for editorial assistance; and members of the Kasper lab for helpful discussions. Electron microscopy was performed at the Harvard Medical School EM Facility with assistance from Louise Trakimas and Maria Ericsson. N.K.S. was supported by NIH K08 AI108690 and a Career Development Award from Boston Children’s Hospital.

## Author contributions

N.K.S. conceived the study, designed and perfomed experiments, and analyzed all data. C.J.C. performed experiments. D.A. generated parabiotic mice. U.H.v.A. helped with data interpretation. D.L.K. supervised all aspects of the project. N.K.S. and D.L.K. wrote the paper. All authors discussed and commented on the manuscript.

## Materials and Methods Mice

### Mice

Timed-pregnant Swiss Webster (SW) mice (Taconic Biosciences), timed-pregnant C57BL/6 mice (Charles River and Jackson Laboratories), C57BL/6 mice (Jackson Laboratories), and B6.SJL-PtprcaPepcb/Boy mice (B6 CD45.1; Charles River and Jackson Laboratories) were maintained under specific pathogen-free conditions. C57BL/6 CD45.1^+^ CD45.2^+^ mice were bred at Harvard University. At E14, pregnant dams were injected intravenously with either 2 mg of an antibody to IL-7Rα (clone A7R34; Bio X Cell; West Lebanon, NH) or an isotype control (clone 2A3; Bio X Cell) to generate PP-deficient and control mice, respectively. For some experiments, pregnant dams were injected with the IL-7Rα antibody on different days ranging between E15 and E18. GF SW mice were bred and maintained in vinyl isolators within the animal facility at Harvard University. For colonization experiments, GF SW mice were gavaged with 100 μl of fecal extract prepared by homogenizing ∼100 mg of feces from control or PP-deficient mice in 1 ml of PBS and filtering the mixture through a 100-μm cell strainer to remove particulate matter. Mice used in experiments were age-matched (typically 8-12 weeks old) and drawn randomly from the same litter, where feasible. The mice used in Figure 2 were 16-18 weeks old. All procedures were approved by the Harvard Medical Area Standing Committee on Animals and are in accordance with NIH guidelines.

### Bacterial culture of MLNs

The middle segments of MLNs, which drain only the jejunum and the ileum^30^, were harvested from control and PP-deficient mice, pressed through a 70-μm cell strainer, and resuspended in 1 ml of PBS. Serial dilutions were plated on both trypticase soy agar with 5% sheep blood (BD Biosciences) and *Brucella* agar with 5% sheep blood, hemin, and vitamin K_1_ (BD Biosciences). Plates were incubated at 37°C in aerobic and anaerobic conditions (80% N_2_, 10% H_2_, 10% CO_2_), respectively, and colonies were counted after 24 h (aerobic) or 48 h (anaerobic).

### Bacterial infections

*Salmonella* infections: *S. enterica* serovar Typhimurium strain SL1344 was grown overnight at 37°C in Luria-Bertani (LB) broth containing streptomycin (200 μg/ml). Control and PP-deficient C57BL/6 mice (12-15 weeks old) were gavaged with ~10^8^ CFU and sacrificed 24 h later. The distal 10 cm of the small intestine was harvested, flushed with PBS, and homogenized with a bead beater and 3.2-mm steel beads. Serial dilutions of the tissue homogenate were plated on LB/streptomycin agar, and colonies were counted after 24 h of incubation at 37°C.

*C. rodentium* infections: *C. rodentium* strain DBS100 (a kind gift from Lynn Bry, Brigham and Women’s Hospital, Boston, MA) was grown overnight in LB broth. Control and PP-deficient C57BL/6 mice (16-20 weeks old) were gavaged with ∼5 × 10^8^ CFU. Fecal samples were collected periodically. The mice were sacrificed 8 days after infection, at which time the proximal 5 cm of colon was harvested and homogenized by bead beating. Serial dilutions of fecal and colonic homogenates were plated on MacConkey agar (BD). Colonies were counted after 24 h of incubation at 37°C; *C. rodentium* colonies were identified by their deep pink, slightly translucent appearance. As a control, fecal samples from uninfected animals were plated on MacConkey agar to ensure that there were no commensal organisms with a similar morphologic appearance.

### Transmission electron microscopy

A 4- to 5-mm length of ileum containing a PP was excised from a control mouse, and three separate 4- to 5-mm sections of ileum (∼1 cm apart) were removed from PP-deficient mice. Two mice per group were examined. All tissue samples were incubated overnight in fixative (1.25% formaldehyde, 2.5% glutaraldehyde, and 0.03% picric acid in 0.1 M sodium cacodylate buffer, pH 7.4). Samples then underwent osmication, uranyl acetate staining, and dehydration in alcohols and were embedded in TAAB 812 Resin (Marivac Ltd.; Nova Scotia, Canada). Sections (80 nm) were cut with an <u>U</u>ltracut microtome (Leica; Buffalo Grove, IL), picked up on 100-mesh formvar/carbon-coated grids, and stained with 0.2% lead citrate. Microscopy was performed with a 1200EX transmission electron microscope (JEOL; Peabody, MA) at 80 kV with primary magnification of 500x–15,000x.

### Intestinal permeability

Control and PP-deficient mice were gavaged with 8 mg of FITC-dextran (4 kDa; Sigma) per 10 g of body weight. The mice were sacrificed 4 h later, and blood was obtained via cardiac puncture. The serum concentration of FITC-dextran was measured with a Synergy HT fluorimeter (BioTek; Winooski, VT) and determined based on an 8-point standard curve.

### Cultured intestinal segments

The distal 5 cm of the ileum and the proximal 5 cm of the colon were harvested, flushed of contents, opened longitudinally, and cultured for 24 h in RPMI containing penicillin/streptomycin and 5% FBS. The supernatant was stored at −80°C until analysis. Concentrations of CCL25 and CCL28 were measured in the supernatant by ELISA (R&D Systems; Minneapolis, MN).

### IgA ELISA

Feces were collected, homogenized in 1 ml of PBS, and filtered through a 100-μm cell strainer to remove particulate matter. For small-intestinal contents, the distal 5 cm of the ileum was harvested and flushed with PBS, and the contents were filtered through a 100-μm cell strainer. The concentrations of IgA in the fecal extract, small-intestinal content extract, and supernatants from the cultured intestinal segments were measured with an IgA ELISA Quantitation Set (Bethyl Laboratories; Montgomery, TX) according to the manufacturer’s instructions.

### Isolation of lymphocytes and flow cytometry

Small-intestinal, cecal, and colonic LP lymphocytes were isolated as previously described^31^. In brief, tissues were collected, cleaned of mesenteric fat, and flushed of intestinal contents, and PPs were removed. The tissue segments were inverted, and the epithelial layer was dissociated by incubation in RPMI containing 1.6% FBS, 0.015% DTT, and 1 mM EDTA for 15 min at 37°C, with constant stirring at 575 rpm. After washing in RPMI to remove residual mucus, the intestinal segments were finely minced and then incubated in RPMI containing dispase (0.5 mg/ml; Sigma), collagenase II (1.5 mg/ml; Sigma), and 1.2% FBS for 40 min at 37°C, with constant stirring at 575 rpm. The digested tissue was serially filtered through a 100-μm and a 40-μm cell strainer prior to flow cytometric analysis. To isolate lymphocytes from spleen, MLNs, and PPs, the organs were pressed through a 70-μm cell strainer. Splenic cells were subsequently incubated in ACK lysis buffer for 2 min on ice and washed in RPMI containing 2% FBS. Lungs were minced and incubated in RPMI containing collagenase (1.5 mg/ml) and DNase I (150 μg/ml; Worthington Biochemical Corp; Lakewood, NJ) for 1 h at 37°C, with constant stirring at 575 rpm. The digested tissue was filtered through a 40-μm cell strainer, incubated in ACK lysis buffer for 2 min on ice, and washed with RPMI containing 2% FBS. For peritoneal lymphocytes, the peritoneal cavity was lavaged with 10 ml of PBS. All samples were resuspended in RPMI containing 2% FBS prior to antibody staining for flow cytometric characterization.

For flow cytometric detection of M cells, small-intestinal epithelial cells were isolated by the above approach. A primary antibody to GP2 (2F11-C3; MBL Life Science), followed by a PE-conjugated secondary antibody (Biolegend), was used.

Fluorophore-conjugated antibodies to the following antigens were used (with clones listed in parentheses): CD45 (30-F11), TCRβ (H57-97), TCRγδ (UC7-13D5), CD4 (GK1.5), CD19 (6D5), CD11b (M1/70), CD11c (N418), B220 (RA3-6B2), IgA (C10-3; BD Biosciences), CD1d (1B1), CD5 (53-7.3), and CD43 (S11). All antibodies were from Biolegend unless indicated otherwise. Cells were permeabilized with the Foxp3/Transcription Factor Staining Buffer Set (eBiosciences) for intracellular staining of IgA. A fixable viability dye conjugated to eFluor 780 (eBioscience) was included in all stains. Flow cytometry was performed with a MACSQuant Analyzer (Miltenyi Biotec), and data were analyzed with FlowJo software (FlowJo; Ashland, OR).

### Adoptive transfer

PPs and spleens were harvested from B6 CD45.1^+^ and B6 CD45.1^+^ CD45.2^+^ mice, respectively. Cells were enumerated, washed twice in PBS, and pooled together. Approximately 3 × 10^6^ PP cells and 1.7 × 10^7^ splenocytes were injected IV into each recipient PP-deficient mouse (C57BL/6). Three days after adoptive transfer, the recipient mice were sacrificed, and the small-intestinal and colonic LPs were assessed by flow cytometry. The mice had ∼3-5% donor cells (either PP cells or splenocytes) in each organ. For CD19^+^ and CD4^+^ cells, we calculated the percentage of PP-derived donor cells as follows: # CD45.1^+^ CD45.2^−^/ # CD45.1^+^ CD45.2^+/−^ [alternatively described as PP cells/(PP cells + splenocytes)].

### Real-time RT-PCR

In accordance with the manufacturers’ instructions, RNA was extracted from ileal and colonic samples with TRIzol (Ambion), and complementary DNA was generated with the SuperScript III First-Strand Synthesis System (Invitrogen). Real-time RT-PCR was performed with a SYBR Select Master Mix (Applied Biosystems) and a LightCycler 480 II (Roche). Values were normalized to the expression of GAPDH for each sample. The following previously validated primer sets were used: GAPDH^32^—F, 5’-CCTCGTCCCGTAGACAAAATG-3’, and R, 5’-TCTCCACTTTGCCACTGCAA-3’; MADCAM1^33^—F, 5’-CCTGAGTCTGAGGTAGCTGTGG-3’, and R, 5’-GAGTGCCTGTGTGTCTGACAGCAT-3’; CXCL13^34^—F, 5’-CAGAATGAGGCTCAGCACAGC-3’, and R, 5’-CAGAATACCGTGGCCTGGAG-3’.

### Parabiosis

Parabiosis surgery was performed as previously described^35^. In brief, sex-matched congenic partners were anaesthetized with a mixture of ketamine (100 mg/kg of body weight), xylazine (10 mg/kg of body weight), and acepromazine (3 mg/kg of body weight) by IP injection. The corresponding lateral aspects of mice were shaved, two matching skin incisions were made from the olecranon to the knee joint of each mouse, and the subcutaneous fascia was bluntly dissected to create about 0.5 cm of free skin. The olecranon and knee joints were attached by a double 4-0 suture, and staples or a continuous 6-0 suture was used to approximate the dorsal and ventral skin flaps. The mice were then kept on heating pads and continuously monitored until recovery. Flunixin (2.5 mg/kg body of weight) and buprenorphine (0.05–0.1 mg/kg of body weight) were administered by subcutaneous injection for postsurgical analgesic treatment. After 2 weeks, chimerism of leukocytes in the spleen was monitored to ensure equivalent blood exchange between parabiotic partners. Parabiotic pairs consisted of PP-deficient mice and WT mice and—as a control—two congenic WT mice.

### TNBS colitis

TNBS colitis was induced as previously described^36^. In brief, SW mice were sensitized on the back with 150 μl of 1% TNBS (Sigma). A week later, the mice were rectally challenged with 60 μg of TNBS (prepared in 50% ethanol)/g of body weight. Daily weights were assessed, and the mice were sacrificed 3 days after rectal challenge. The disease score represents the summation of the following features: (1) weight loss (scored as follows: <5%=0, 5–10%=1, 10–15%=2, 15–20%=3, and >20%=4); (2) colon length (scored as follows: >9 cm=0, 8–9 cm=1, 7–8 cm=2, 6–7 cm=3, <6 cm=4); (3) stool consistency (scored as follows: normal appearing=0, soft pellets=1, soft stool with no formed pellets=2, liquid stool=3, appearance of frank blood=4); (4) colon thickness (subjectively scored as 0–4, with 0 being normal); and (5) histology (scored as 0–4, with 0 being normal, by a pathologist blinded to treatment group). As a control, mice that were not sensitized with TNBS and rectally challenged with 50% ethanol (without TNBS) had disease scores of 0–2.

### Quantitative rDNA analysis

To quantitatively assess the burden of mucosa-associated bacteria in the intestine, the distal 5 cm of ileum or colon was harvested and flushed with PBS to remove luminal bacteria. The remaining tissue was opened longitudinally and vigorously washed in PBS to liberate the bacteria more closely associated with the epithelium and mucus layer. Genomic DNA was isolated from these mucosa-associated bacterial populations by bead beating followed by phenol-chloroform extraction; fluorescent primer- and probe-based chemistry and a LightCycler 480 II (Roche) were used to quantitate 16S rDNA as previously described^37^.

### Statistics

Sample size estimates for each experiment were based on prior lab experience. Prism 6 (GraphPad Software; La Jolla, CA) was used for all statistical analyses. All *p* values were calculated by unpaired, two-tailed Student’s *t* or Mann-Whitney test, as appropriate. Data from the parabiosis experiment were analyzed by Wilcoxon test. In dot plots shown in the figures, each data point represents an individual mouse, and horizontal bars represent the median value.

