## Extended Data Figures for "Anatomically remote education of B cells is required for colonic health"

### Extended Data Figure 1

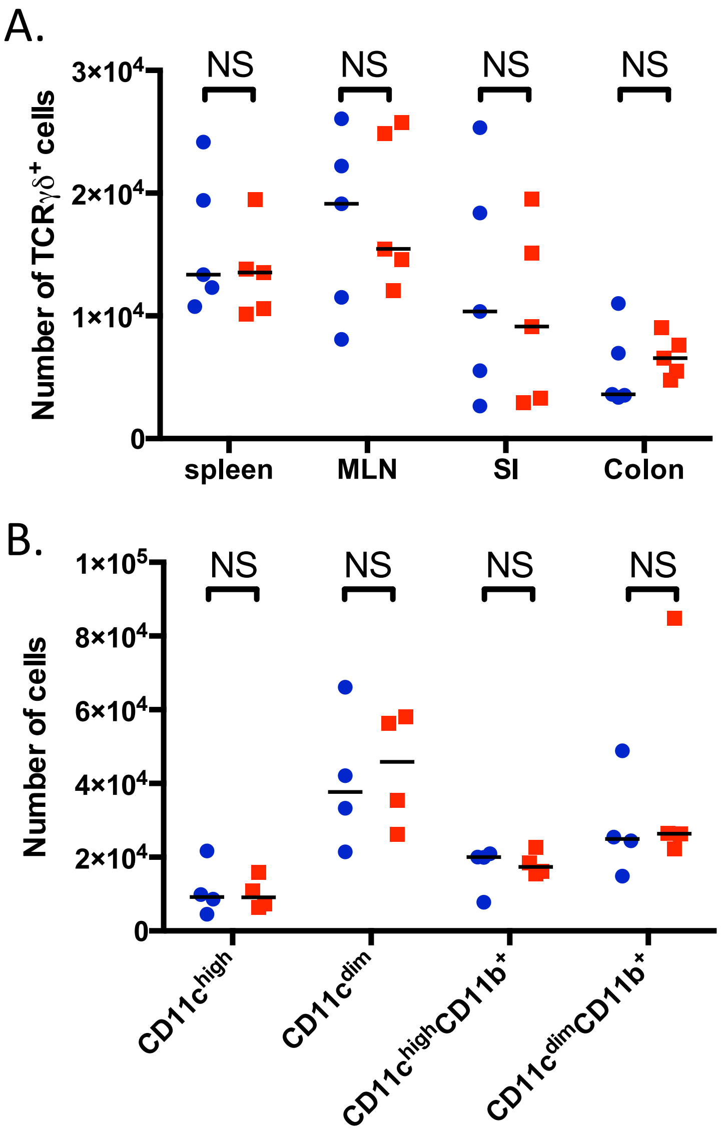


**Extended Data Figure 1. PP-deficient mice have no defect in TCRγδ+ or myeloid-lineage cells.** A. Numbers of TCRγδ+ cells in the spleen, MLNs, small intestine (SI), and colon. B. Numbers of various myeloid cell populations present in the colon. Control animals are shown in blue circles, PP-deficient animals are depicted by red squares. NS, not significant.

### Extended Data Figure 2

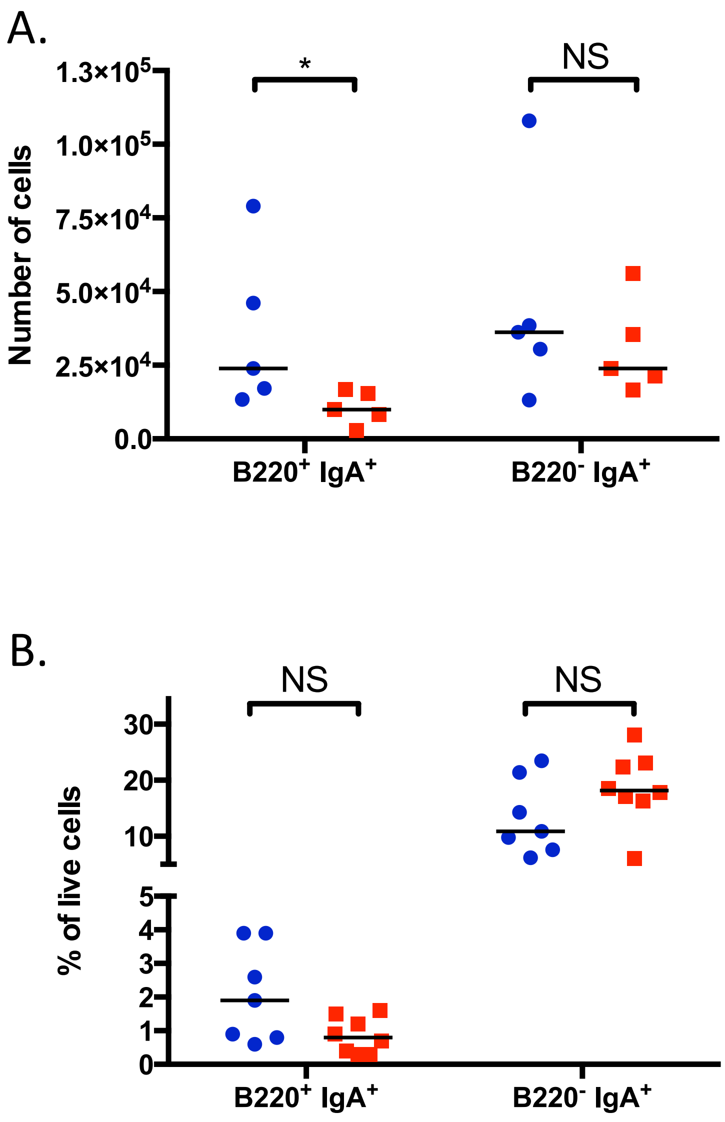


**Extended Data Figure 2. PP-deficient mice have reduced numbers of colonic plasmablasts with no change in small-intestinal frequencies.** A. Numbers of plasmablasts ($B220^+$ $IgA^+$) and plasma cells ($B220^-$ $IgA^+$) in the colon. B. Frequency of plasmablasts and plasma cells in the small-intestinal lamina propria. Control animals are shown in blue circles, PP-deficient animals are depicted by red squares. NS, not significant; *, $p<0.05$.

### Extended Data Figure 3

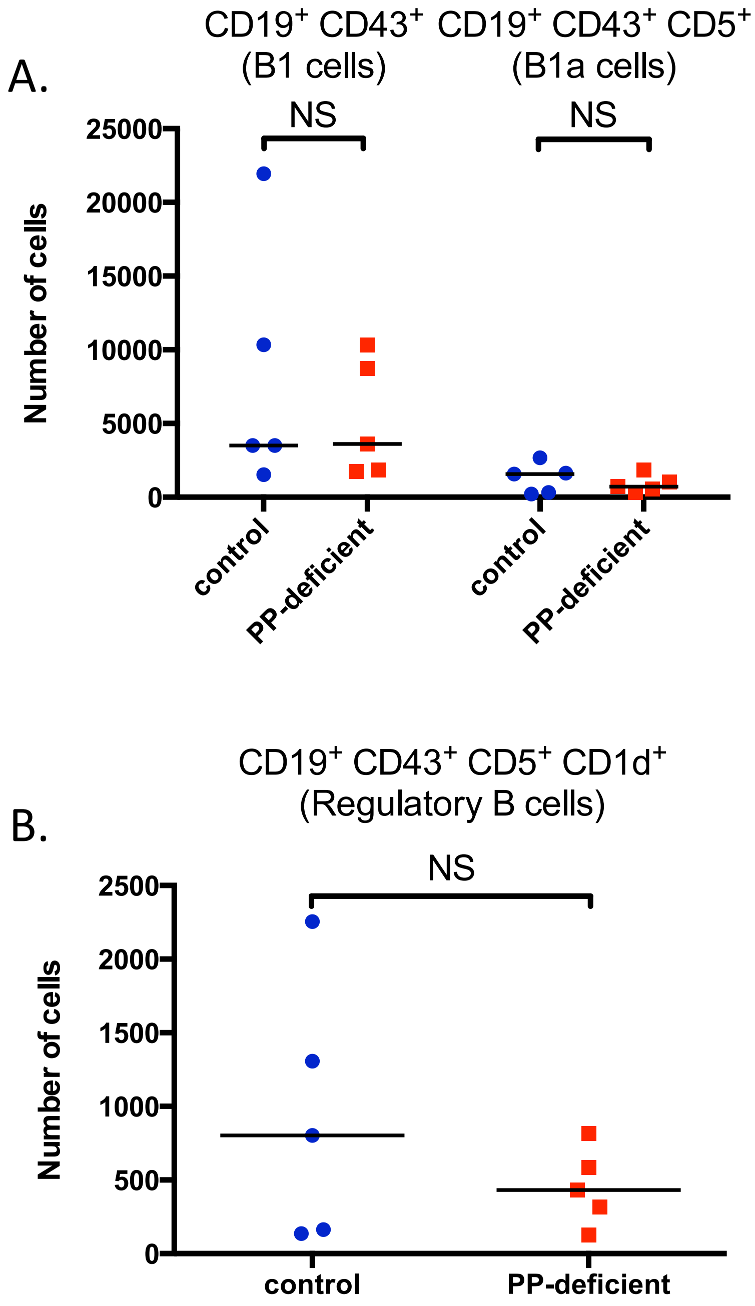


**Extended Data Figure 3. PP-deficient mice have no defect in colonic B1 cells**. Numbers of B1 cells ($CD19^+$ $CD43^+$), B1a cells ($CD19^+$ $CD43^+$ $CD5^+$) (A), and regulatory B cells ($CD19^+$ $CD43^+$ $CD5^+$ $CD1d^+$) (B) in the colon. NS, not significant.

### Extended Data Figure 4

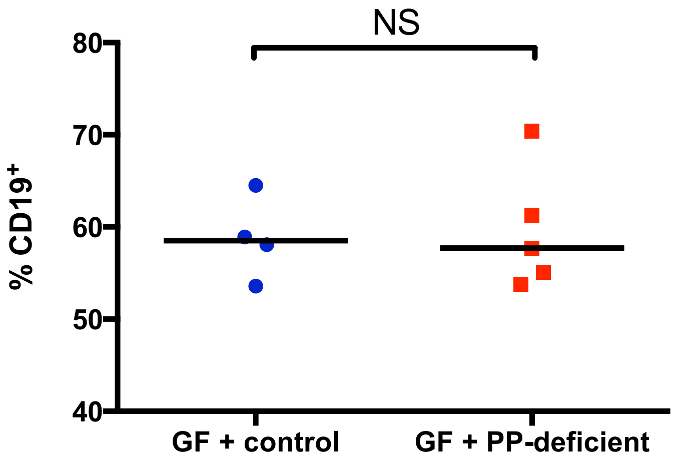


**Extended Data Figure 4. Colonization of GF animals with the microbiota of PP-deficient mice does not affect the colonic frequency of B cells.** Frequency of $CD19^+$ cells (gated on live $CD45^+$ cells) in the colonic lamina propria of gnotobiotic animals colonized with the microbiota of either control or PP-deficient mice. NS, not significant.

Extended Data Figure 5

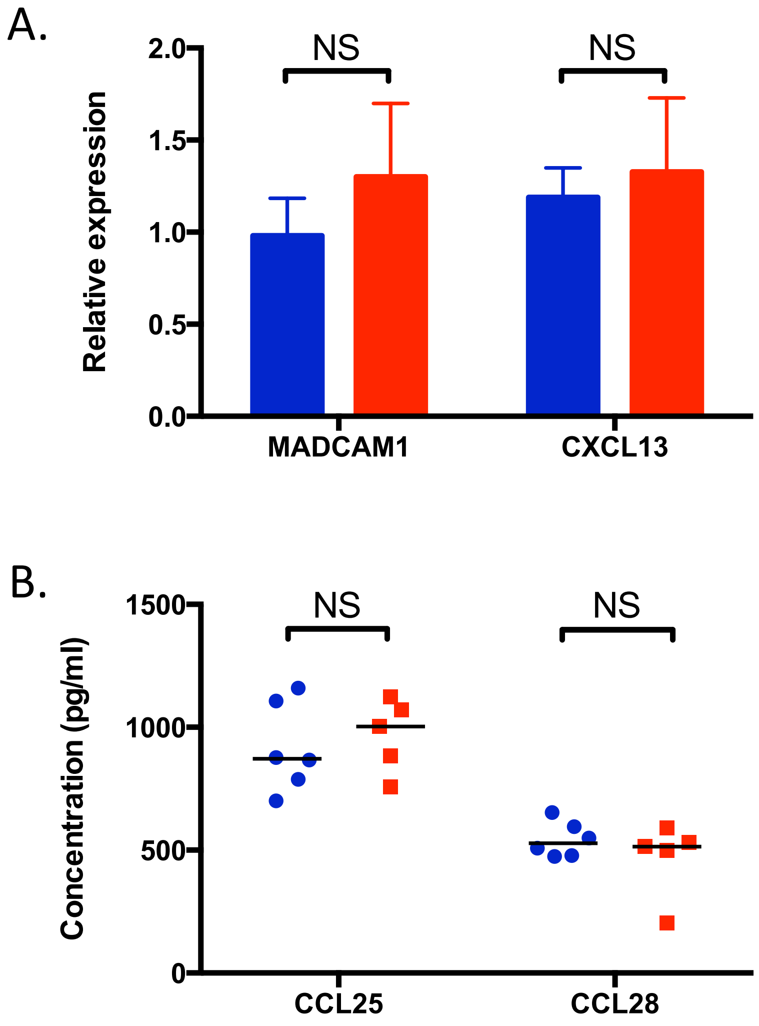


**Extended Data Figure 5. Control and PP-deficient mice do not differ in colonic levels of relevant addressins and chemokines.** A. qPCR analysis of expression levels of colonic MADCAM1 and CXCL13 from control (blue bars) and PP-deficient animals (red bars; *n*=4 per group). B. Concentration of CCL25 and CCL28 in colonic segments from control (blue circles) and PP-deficient mice (red squares). NS, not significant.

### Extended Data Figure 6

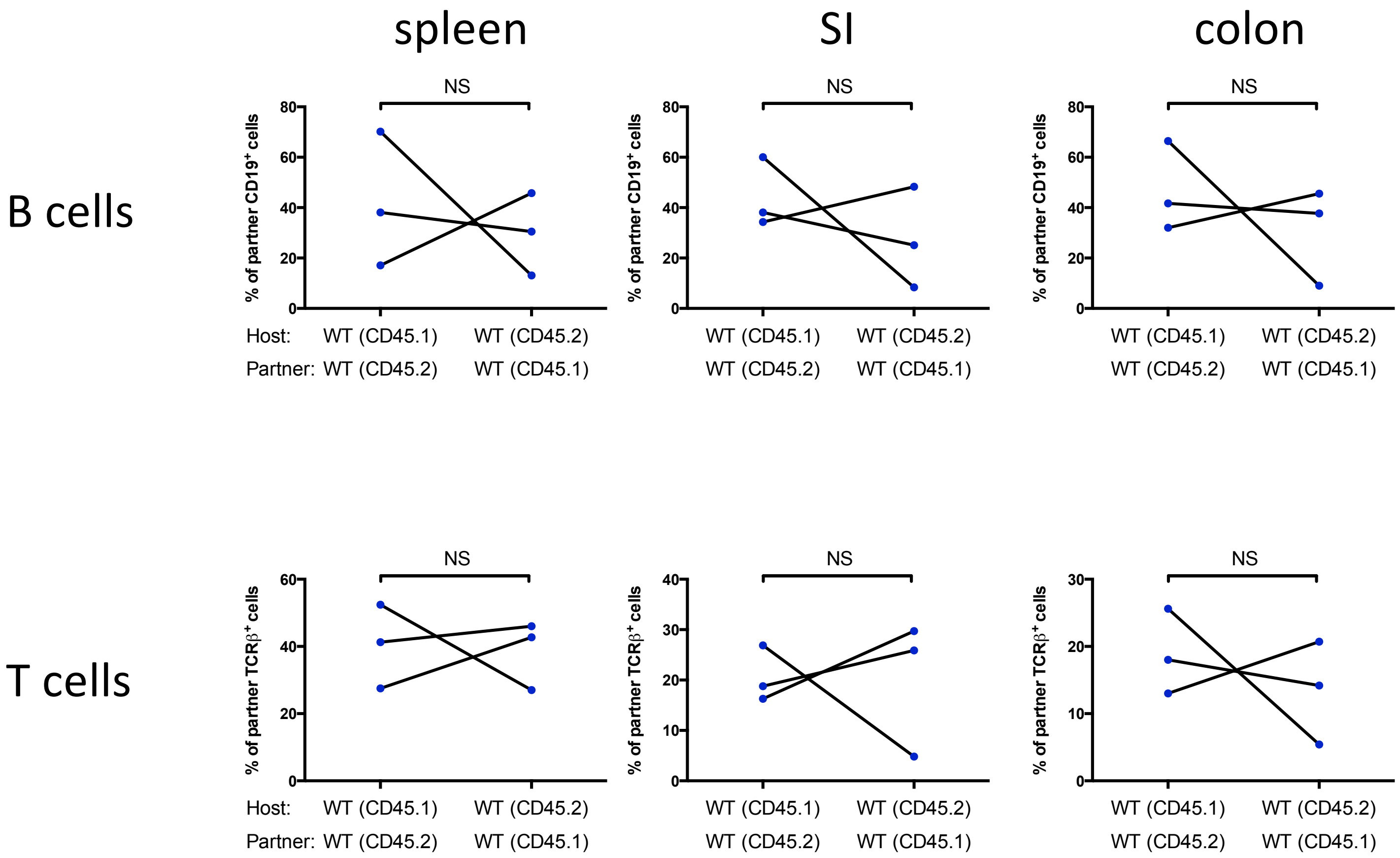


**Extended Data Figure 6. CD45 allotype does not affect recruitment of B and T cells in the spleen and intestines.** Percentage of CD19⁺ cells (top row) or TCRβ⁺ cells (bottom row) in the spleen (left column), small intestine (SI; middle column), and colon (right column) of the indicated host mouse that is derived from the partner. Lines connect parabiont partners.

### Extended Data Figure 7

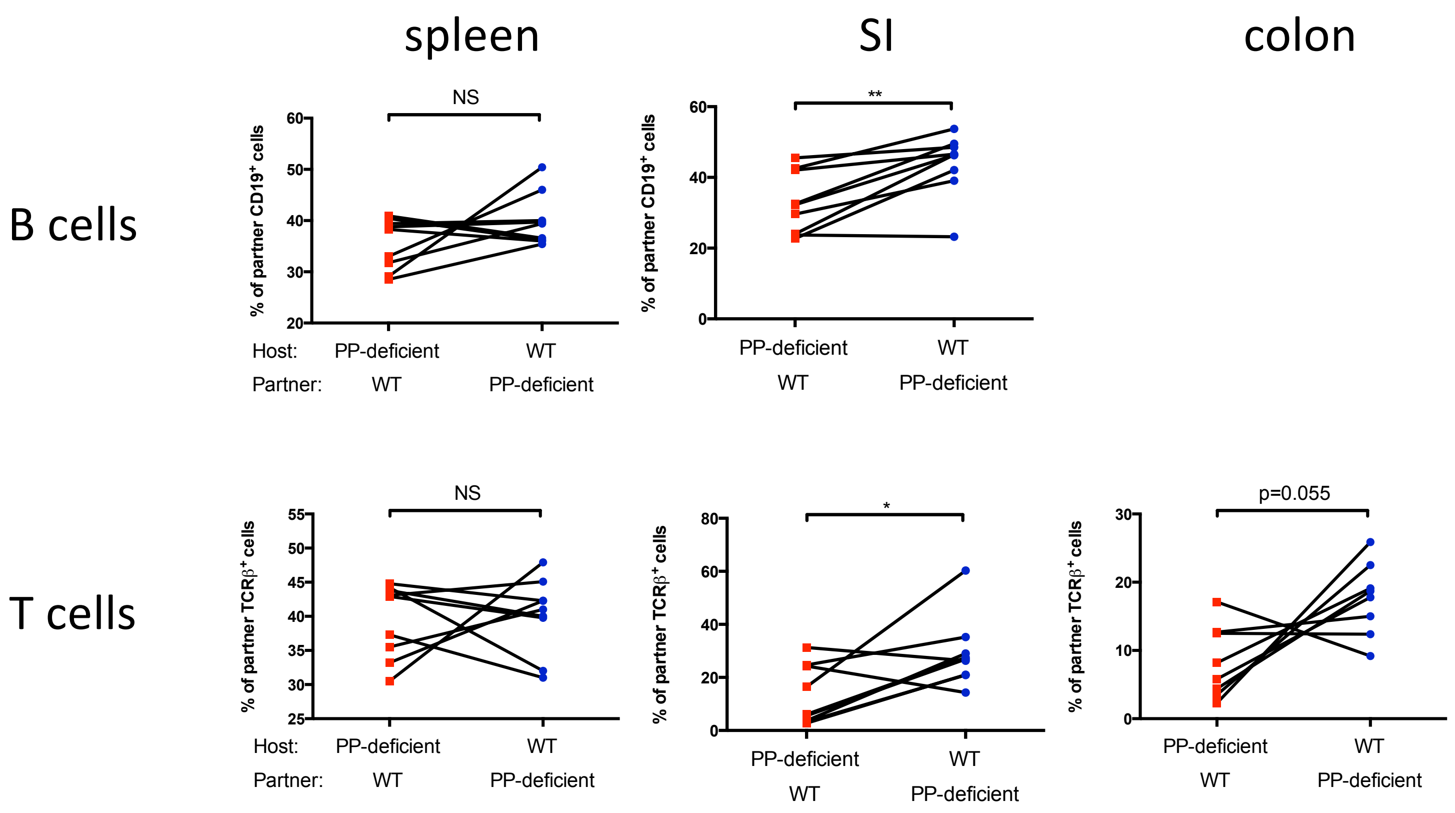


**Extended Data Figure 7. PP-deficient mice have a defect in recruitment of B and T cells to the intestines.** Percentage of $CD19^+$ cells (top row) or $TCR\beta^+$ cells (bottom row) in the spleen (left column), small intestine (SI; middle column), and colon (right column) of the indicated host mouse that is derived from the partner. Lines connect parabiont partners.

### Extended Data Figure 8

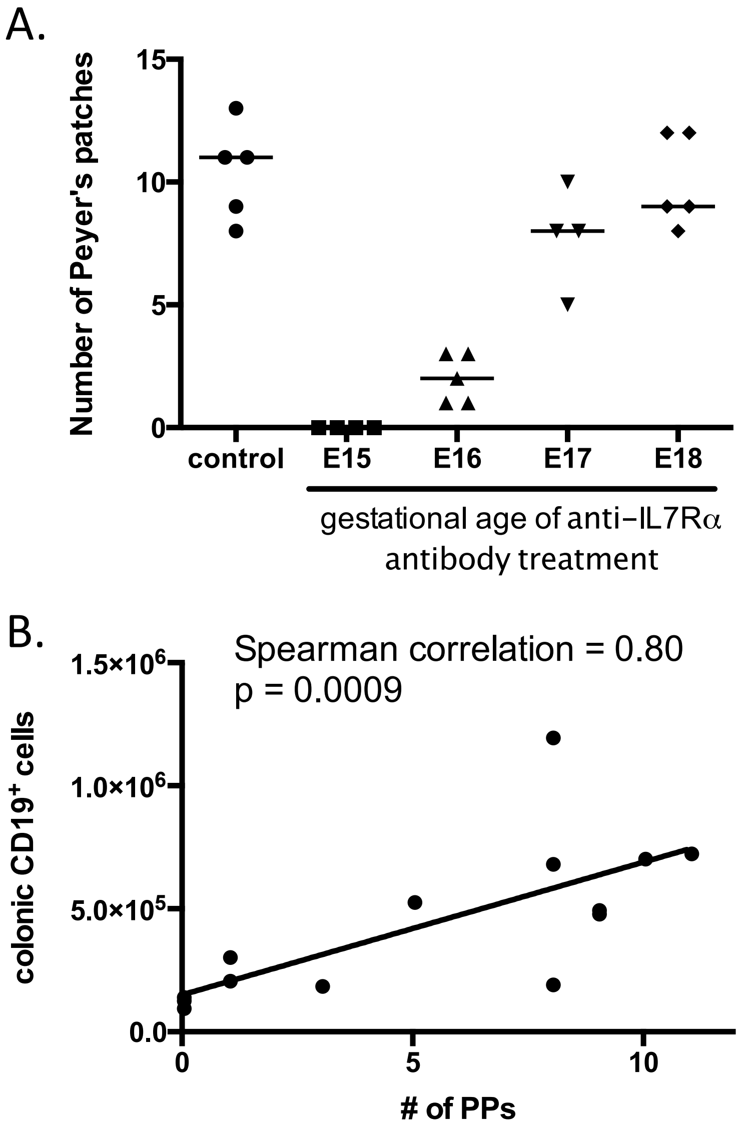


**Extended Data Figure 8. The number of colonic B cells correlates with the number of PPs.** A. Numbers of PPs in control mice and mice exposed to antibody to IL-7Rα at E15, E16, E17, or E18. B. Correlation between the number of colonic $CD19^+$ cells and the number of PPs. *r* value reflects the Spearman correlation.

### Extended Data Figure 9

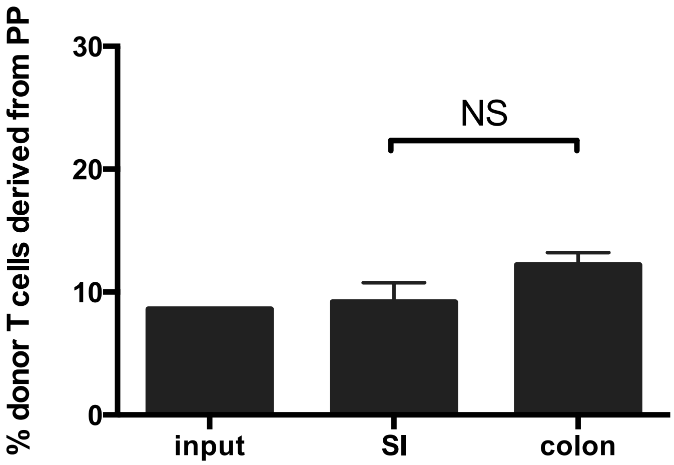


**Extended Data Figure 9. PP-derived T cells do not preferentially migrate to the colon.** Shown are the fractions of donor T cells that are PP-derived in the input, the small intestine (SI), and the colon.

### Extended Data Figure 10

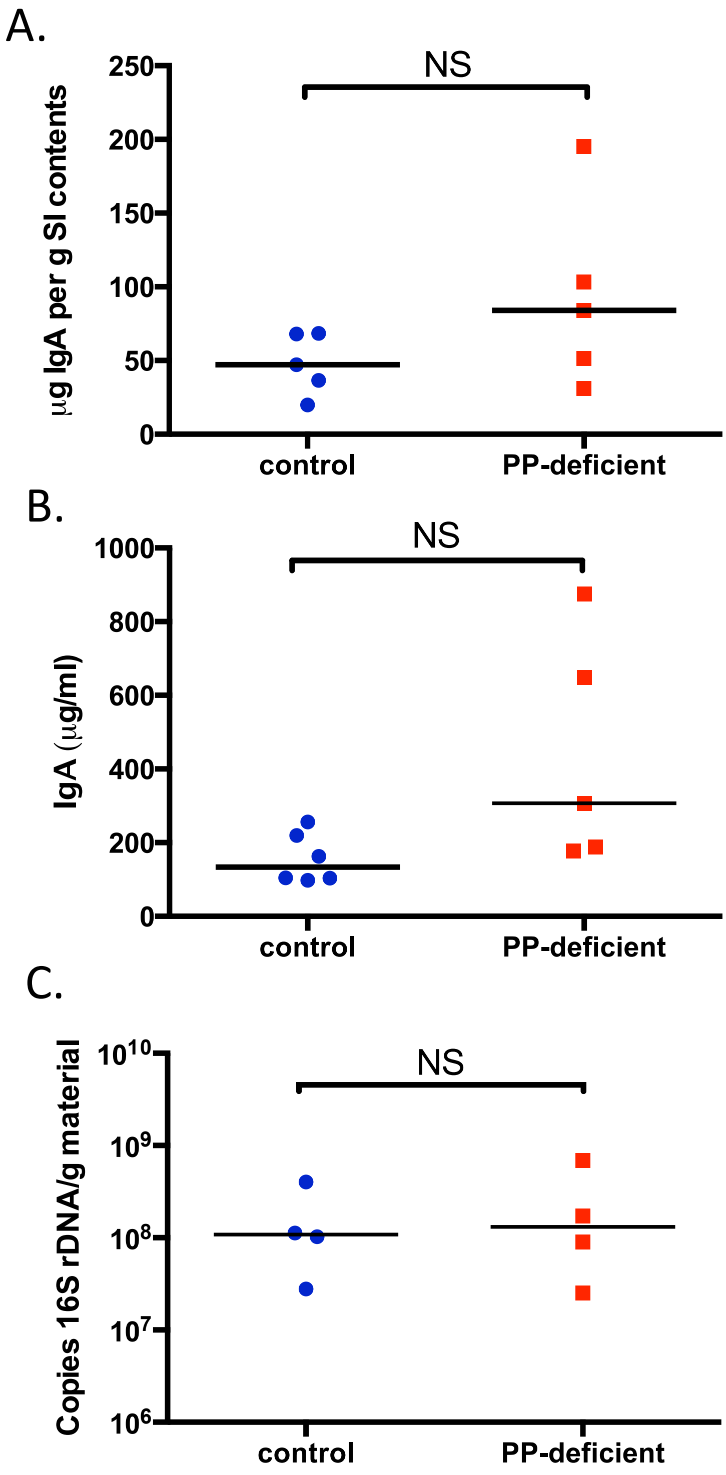


**Extended Data Figure 10. PP-deficient mice have no defect in IgA production or level of mucosa-associated bacteria in the small intestine.** A. Levels of IgA in distal ileal contents. B. Ileal segments were harvested from control and PP-deficient mice and cultured *ex vivo*. The IgA concentration in the supernatant was measured. C. qPCR-based quantification of mucosa-associated bacteria in the small intestine is shown. NS, not significant. Data are representative of ≥2 experiments.
